# The interaction between abiotic and biotic soil factors drive heterosis expression in maize

**DOI:** 10.1101/2024.08.30.610574

**Authors:** Kayla M. Clouse, Martel L. Ellis, Natalie E. Ford, Rachel Hostetler, Peter J. Balint-Kurti, Manuel Kleiner, Maggie R. Wagner

**Affiliations:** Department of Ecology and Evolutionary Biology, University of Kansas, Lawrence, KS 66045; Kansas Biological Survey & Center for Ecological Research, University of Kansas, Lawrence, KS 66045; Department of Plant Science, Pennsylvania State University, University Park, PA 16802; Department of Entomology and Plant Pathology, North Carolina State University, Raleigh, NC 27695; Plant Science Research Unit, Agricultural Research Service, United States Department of Agriculture, Raleigh, NC 27695; Department of Plant and Microbial Biology, North Carolina State University, Raleigh, NC 27695

## Abstract

Heterosis or hybrid vigor refers to the superior phenotypes of hybrids relative to their parental inbred lines. Recently, soil microbes were identified as an environmental driver of maize heterosis. While manipulation of the soil microbial community consistently altered heterosis, the direction of the effect appeared to be dependent on the microbiome composition, environment, or both. Abiotic factors are well-known modifiers of heterosis expression, however, how the interactive effects between the soil microbial community and abiotic factors contribute to heterosis are poorly understood. To disentangle the proposed mechanisms by which microbes influence heterosis, we characterize the variation in heterosis expression when maize was grown in soil inocula derived from active maize farms or prairies. While we did not observe consistent differences in heterosis among plants grown in these inocula, our observations reaffirm that microbial effects on heterosis are likely specific to the local microbial community. The introduction of a nutrient amendment resulted in greater heterosis expression in the presence of an agricultural inoculum but not a prairie inoculum. We also observed an effect of soil inocula and nutrient treatment on the composition of bacterial and fungal communities in the root endosphere. In addition, the interaction between soil and nutrient treatment significantly affected bacterial community composition, whereas fungal community composition was only marginally affected by this interaction. These results further suggest that the soil microbial community plays a role in maize heterosis expression but that the abiotic environment is likely a larger driver.

## Introduction

The plant microbiome is composed of bacteria, fungi, archaea, and protists that exist on and within plant compartments. These microorganisms can provide numerous benefits to their hosts, including protection against pathogens (van Wees et al., 2008), increased tolerance to drought (Rolli et al., 2014), nutrient acquisition (Reed et al. 2011), and enhanced plant productivity (Compant et al., 2010). The plant microbiome is strongly structured by both abiotic and biotic soil factors (Fierer et al., 2017), which can in turn affect plant performance. In agricultural systems, conventional management practices such as pesticide application (Walder et al., 2022), tillage (Kraut-Cohen et al., 2020), and fertilization (Bahulikar et al., 2019) can lead to shifts in soil properties and microbial community composition. Furthermore, conventional monoculture cropping practices decrease soil microbiome diversity (Li et al. 2019) and enrich plant pathogens (Zhou et al. 2017).

Plants are similarly strong drivers of microbial communities, assembling microbiomes that are taxonomically and functionally distinct from the soil. Plants recruit microbiome members from the soil through changes in morphology (Oldroyd 2013), immune response (Lebeis et al., 2015), and root exudation (Sasse et al., 2018). For example, plants may modulate their immune system to protect against pathogens through the enrichment of beneficial microbes (Liu et al 2020). Similarly, root exudates from different plant species can stimulate or suppress soil bacteria to select for specific rhizosphere bacteria (Dhungana et al., 2023). Genotypes of the same plant species also vary in their recruitment mechanisms, which can result in distinct root and rhizosphere communities from the same environment (Singer et al. 2019).

The fact that genetically-controlled plant traits shape microbiome assembly suggests that microbiome properties can be inherited. If the underlying genes have mostly additive effects, then the plants’ microbiomes would be expected to be intermediate to those of their parents. However, this is contradicted by data from field-grown maize, in which hybridization between plant genotypes results in distinct bacterial and fungal rhizosphere communities in hybrid relative to parental genotypes (Wagner et al. 2020). Hybrid maize is also more likely to be colonized by beneficial arbuscular mycorrhizal fungi, as well as nitrogen-fixing bacteria (Picard et al. 2008) than inbred maize. *Pseudomonas* strains that produce the beneficial antifungal compound *phlD* are also more abundant in the rhizosphere of hybrid maize (Picard et al. 2004). Together, these observations indicate that microbiome composition, especially the abundance of symbiotic microbes, differ between inbred and hybrid maize.

The deviation of maize microbiome composition from the mid-parent expectation is typical of many other maize phenotypes, including height and yield. This phenomenon, known as heterosis or hybrid vigor, typically refers to the superior phenotypes of hybrid plants relative to their parental inbred lines. Heterosis can vary substantially depending on the plant trait of interest and the environment. The majority of studies exploring environmental effects on heterosis have employed abiotic stress conditions (Li et al., 2022). In these studies, the hybrid genotypes are generally less variable under stress conditions than the inbred genotypes (Knight 1973). However, we have a limited understanding of heterosis expression under normal field conditions.

Recently, the soil microbial community has been found to influence the expression of heterosis for traits such as root biomass in maize (Wagner et al. 2021). In three separate experiments, the elimination or reduction of soil microbes resulted in weakened heterosis, which was due to reduced performance of the inbreds rather than increased performance of the hybrid. Two possible explanations for this observation are that (1) hybrids may be more resistant than inbreds to pathogenic soil microbes, or (2) inbreds but not hybrids mount costly defense responses to non-pathogenic microbes. However, these hypothesized mechanisms are not consistent with the results of a fourth experiment conducted in a separate environment, in which the reduction of soil microbes resulted in greater heterosis. This indicates that the exact role of microbial communities in heterosis expression depends on the microbiome composition, the abiotic environment, or both. It also suggests alternative mechanisms of microbe-dependent heterosis that involve interactions with growth-promoting organisms, such as: (3) hybrids may host greater numbers of beneficial microbes than inbreds do, or (4) inbreds may be more reliant on nutrient-providing microbes than hybrids are. In these scenarios, resource availability in the soil is likely to modify the relationships between the microbiome and host phenotype.

To disentangle the proposed mechanisms by which microbes influence maize heterosis, we characterized the variation in better parent heterosis (BPH), herein referred to as heterosis, when inbred and hybrid maize were grown in soil inocula derived from active maize farms or prairies. Due to recent monoculture cropping, we expected a greater abundance of pathogens, as well as the loss of disease-suppressive functional groups, in the agricultural inocula relative to the prairie inocula. Resultantly, if microbial effects on heterosis are due to soil pathogens then we expect that heterosis will be stronger in the agricultural inocula due to decreased performance of the inbred genotypes. Furthermore, to test the interactive effects of soil inocula and the environment, we introduced a nutrient amendment. We expected the nutrient treatment to decrease the abundance and diversity of symbiotic microbes in roots, since the plants will be less reliant on them for nutrient acquisition. Finally, if inbreds are more dependent than hybrids on nutrient-providing organisms, then we would expect to see a weaker effect of inocula on heterosis in the high nutrient treatment.

## Methods

### Experiment 1

To characterize heterosis in response to soil inoculum source, we collected four soils from agricultural and prairie fields in eastern Kansas. The two agricultural soils were collected from maize farms (Lawrence, KS), which have been in maize production for over 80 years, and the two prairie soils were collected from Welda Prairie (Welda, KS) and Clinton Wildlife Reserve (Tecumseh, KS). We also included a “control” soil, which contained an equal ratio of each soil and was steam-sterilized. The soils were used as inocula for two inbred genotypes of maize (B73 and Mo17) and their F1 hybrid (B73xMo17). Prior to planting, the seeds were surface-sterilized for three minutes with 5% sodium hypochlorite followed by 70% ethanol then rinsed with sterile distilled water three times. The surface-sterilized seeds were air-dried in a biosafety cabinet then two seeds per genotype were planted in cone-tainer pots (SC7R; Stuewe & Sons) containing a mix of sterile calcined clay (“Pro’s Choice Rapid Dry”; Oil-Dri Corporation) and sterile potting soil. The soil inoculum (15% v/v total soil) was added on top of the seeds followed by additional sterile calcined clay until each pot was full. Fifteen plants per genotype per soil inocula (N=225) were placed in eight randomized blocks (28 plants/block) in a growth chamber (12-hr days, 27℃/23℃, ambient humidity) then 45 mL of sterile 0.25x Murashige and Skoog (MS) nutrient solution was added to each pot. The plants were grown for four weeks and watered approximately every two days with UV-sterilized water. Emergence was measured daily for the first 10 days of the experiment. After one week of growth, the plants were thinned to one seedling per pot. Plant height was measured weekly throughout the experiment and chlorophyll content was recorded at two and four weeks using the MC-100 Chlorophyll Concentration Meter (Apogee Instruments). Three chlorophyll measurements were recorded approximately 5 cm from the leaf tip of the uppermost collared leaf and then the three measurements were averaged. Stem diameter was measured once at four weeks using a digital caliper (Mitutoyo Corp.) between the first and second emerged leaf, about 4 cm above the base of the stem. After four weeks of growth, the roots were separated from the shoots then dried for 48 hours at 80°C for biomass measurements.

### Experiment 2

To characterize heterosis in response to nutrient amendments, we introduced a high-nutrient or low-nutrient solution to a subset of the same soil inocula used in Experiment 1. The soil inocula that induced the highest (prairie soil 1) and lowest (agriculture soil 1) average heterosis across plant traits were selected for this experiment. Two seeds of the same genotypes and source from Experiment 1 (B73, Mo17, and B73xMo17) were surface-sterilized then planted in cone-tainer pots containing a mix of sterile calcined clay and potting soil. The agricultural or prairie soil inoculum (15% v/v total soil) was then added on top of the seeds followed by additional sterile calcined clay. Fifteen plants per genotype per soil inocula per nutrient treatment (N=180) were placed in seven randomized blocks (28 plants/block) in a growth chamber (12-hr days, 27℃/23℃, ambient humidity) then 45 mL of either sterile 1x or 0.1x Hoagland’s No. 2 Basal Salt Mixture (Caisson Laboratories, Inc.) was added to each pot. The plants were watered every two days for four weeks with UV-sterilized water and at every other watering the plants received 45 mL of sterile 1x (high nutrient) or 0.1x (low nutrient) Hoagland’s solution in lieu of water. Emergence was measured daily for the first 10 days of the experiment. After one week of growth, the plants were thinned to one seedling per pot. Plant height, chlorophyll content, and stem diameter were measured at two and four weeks as described in Experiment 1. Soil pH was measured using a pH probe (Hanna Instruments) at two and four weeks. After four weeks of growth, the plants were uprooted then 2.5 cm fragments from the bottom of the primary root were collected from five plants per genotype per soil per treatment. The remaining roots were separated from the shoots then oven-dried for biomass measurements.

### Total leaf nitrogen and carbon

For Experiment 2, we ground dried shoot tissue from three plants per genotype per soil inoculum per treatment using a mortar and pestle. The coarse tissue was transferred to 2 mL tubes containing 4 6-mm metal beads then homogenized at 1400 rpm for 5 minutes using the Ohaus HT Homogenizer (Ohaus Corporation). The homogenized tissue was transferred to coin envelopes then placed in a drying oven at 55°C for one week. After drying, the samples were transferred to a desiccator for 48 hours then two technical replicates per sample were weighed (4.7-5.3 mg per replicate) in tin capsules using a microbalance. The tin capsules were folded then stored in a desiccator until processing on the FlashSmart™ Elemental Analyzer (Thermo Scientific™). The analyzer was calibrated using the Acetanilide standard curve then carbon and nitrogen gas concentration was determined using dry combustion followed by gas chromatography.

### Statistical analyses

All data analysis was performed using R version 4.3.2, particularly the packages tidyverse (Wickham et al. 2019), lme4 (Bates et al., 2015), lmerTest (Kuznetsova et al., 2017), emmeans (Lenth 2024), ggpubr (Kassambara 2023) and vegan (Oksanen et al. 2022). For Experiment 1, a two-way ANOVA with Type III sum of squares was applied to linear mixed-effect models for each plant trait with Genotype, Soil Inoculum, and their interaction as fixed predictor variables and Block as a random-intercept term. For Experiment 2, a three-way ANOVA with Type III sum of squares was applied to linear mixed-effect models for each plant trait with Genotype, Soil Inoculum, Treatment, and their interactions as predictor variables and Block as a random-intercept term. Pairwise contrasts between each inbred and the hybrid for each plant trait were performed using Dunnett’s post-hoc procedure. Emergence success was compared between the inbreds and hybrid using Fisher’s exact test. Resulting p-values were adjusted for multiple comparisons (Benjamini and Hochberg 1995) for each experiment.

### Heterosis calculations and statistical inference

For both experiments, the estimated marginal mean was extracted from linear mixed-effect models to calculate better parent (BPH) and mid-parent (MPH) heterosis for each plant trait. To calculate BPH and MPH the following equations were used: BPH = (B73xMo17 - max(B73, Mo17))/(max(B73, Mo17)) and MPH = (B73xMo17 - (B73 + Mo17)/2)/((B73 + Mo17)/2). Next, we calculated “ΔBPH” as the pairwise difference in BPH between agriculture, prairie, and control soils in Experiment 1 and high-nutrient and low-nutrient treatments for each soil in Experiment 2. Positive values of ΔBPH indicated that heterosis was stronger in agriculture versus prairie (and agriculture or prairie versus control) soil or in the high-nutrient versus low-nutrient treatment. Negative values of ΔBPH indicated the reverse. To determine statistical significance of the observed ΔBPH, we recalculated ΔBPH for 999 datasets that were permuted with respect to soil or nutrient treatment to create a distribution of ΔBPH values that would be expected if soil or nutrient treatment had no effect on heterosis. Finally, we compared the observed ΔBPH to the expected null distributions to examine the null hypothesis that heterosis is equally strong in agriculture, prairie, and control soil or high versus low nutrient treatment. To determine the amount of variation in each genotype’s phenotypic response due to soil inocula, we calculated the coefficient of variation for each genotype using the estimated marginal mean for each plant trait. The coefficient of variation was calculated across predictor variables in Experiment 1 (Soil Inoculum) and Experiment 2 (Soil Inoculum and Nutrient Treatment).

### DNA extraction

Root fragments were rinsed with distilled water then placed in cluster tubes with metal beads. Next, roots were freeze-dried and homogenized into a fine powder using the Ohaus HT Homogenizer. The homogenized root tissue was transferred to a 2 mL 96-well plate containing 800 µL of lysis buffer (100 mM NaCl, 10 mM EDTA, 10 mM Tris, pH 8.0) and 1 mm diameter garnet beads (BioSpec, Bartlesville). Next, 10 µL of 20% SDS was added to each well then the plates were homogenized at 20 Hz for 20 min and incubated at 55°C for 90 min. After centrifuging at 4500 x g for 6 min, 400 µL of supernatant was transferred to new 1 mL 96-well plates containing 130 µL of 5 M potassium acetate. Next, the plates were vortexed and incubated at −20°C for 30 min then centrifuged (4500 × g for 6 min). 400 µL of supernatant was transferred to new 1 mL 96-well plates then vortexed with 600 µL of solid phase reversible immobilization bead solution (Rohland and Reich 2012). After allowing the beads to bind to DNA for 10 min, the plates were centrifuged (4500 × g for 6 min) then placed on a magnetic rack for 5 min. The supernatant was removed then the immobilized beads were washed three times with 900 µL of ethanol (80% v/v). After removing the ethanol, the samples were air dried and 75 µL of preheated (37°C) 1× Tris-EDTA buffer (pH 7.5) was added to each well to elute DNA. Finally, the plates were gently vortexed and placed back on a magnet rack then the supernatant was transferred to 0.45 mL 96-well plates.

### PCR and amplicon sequencing

To prepare libraries for 16S-v4 and ITS1 rRNA gene sequencing, we used 5 µL of DreamTaq Hot Start PCR Master Mix (Thermo Scientific) and 2.5 µL of template DNA per reaction. To amplify the 4^th^ variable region of the 16S rRNA gene, we also included 0.4 µL of forward primer (515f), 0.4 µL of reverse primer (806r) (Apprill et al. 2015; Parada et al. 2016), 1.05 µL of PCR-grade water, and 0.15 µL of 100 µM of peptide nucleic acids (PNA) per PCR reaction. The 16S PCR thermocycler settings included a 2 min denaturing cycle at 95°C then 27 cycles of 20 s at 78°C, 5 s at 52°C, and 20 s at 72°C, followed by a 10-min extension at 72°C. To amplify ITS genes, we also included 0.4 µL of forward (ITS1f), 0.4 µL of reverse (ITS2) (Smith and Peay 2014), and 1.7 µL of PCR-grade water per PCR reaction. The ITS1 PCR thermocycler settings included a 2-min denaturing cycle at 95°C then 27 cycles of 20 s at 95°C, 20 s at 50°C, and 50 s at 72°C, followed by a 10-min extension at 72°C. The 16S and ITS PCR product then underwent a second PCR to attach Illumina adapters with indexes. For this PCR, we used 0.8 µL of 10 µM primer mix that contained forward and reverse barcoded primers with P5 and P7 Illumina adaptors. This PCR also included 5 µL of DreamTaq Hot Start PCR Master Mix, 0.15 µL of 100 µM of PNA, and 1 µL of template DNA per reaction. The PCR thermocycler settings included a 2-min denaturing cycle at 95°C then 8 cycles of 20 s at 78°C, 5 s 52°C, and 20 s 72°C, followed by a 10-min extension at 72°C. After the PCR reactions were complete, 10 µL of each 16S-v4 and ITS1 reaction product was pooled. Each pool was normalized using the ‘Just-a-Plate’ kit (Charm Biotech) then DNA was quantified using the Quantus™ fluorometer (Promega). The final pools were combined in equal molarity and sequenced on the Illumina platform Novaseq 6000 at 250 bp PE.

### Sequence processing and quality filtering

Cutadapt (Martin 2011) was used to trim forward and reverse primers from raw sequences before quality filtering. Next, Dada2 (Callahan et al. 2016) was used to remove reads with ambiguous bases or more than two errors. Then the forward and reverse 16S reads were truncated at 235 base pairs. Next, we denoised the reads to classify amplicon sequence variants (ASVs) then removed chimeric ASVs. We used the RDP classifier (Cole et al. 2014) training set 16 and the UNITE database (Nilsson et al., 2019) to assign taxonomy to bacterial and fungal ASVs, respectively. We discarded ASVs that could not be classified at the kingdom level, as well as ASVs that were identified as plant sequences. In addition, we removed samples with less than 300 and 50 usable reads for bacteria and fungi, respectively. In sum, our filtering processes reduced the number of bacterial ASVs from 13831 to 2541 and fungal ASVs from 2170 to 206. However, 97.8% of both the original bacterial and fungal reads were retained after sequencing. Finally, we applied a centered log ratio (CLR) transformation to the final observations in each sample using the ALDEx package (Fernandes et al., 2013).

### Microbiome analysis

All data analysis was performed using R version 4.3.2, particularly the packages tidyverse (Wickham et al. 2019), lme4 (Bates et al., 2015), lmerTest (Kuznetsova et al., 2017), vegan (Oksanen et al. 2022), phyloseq (McMurdie and Holmes, 2014), genefilter (Gentlemen et al. 2023), ALDEx2 (Fernandes et al., 2013), and microViz (Barnett 2024). We used untransformed data to calculate two alpha diversity metrics (Inverse Simpson and Shannon indices). These metrics were modeled using Genotype, Soil Inoculum, Treatment, and their interactions, as well as Sequencing Depth, as fixed predictor variables and Block as a random-intercept term. We assessed these linear mixed-effect models using ANCOVA then adjusted the p-values for multiple comparisons (Benjamini and Hochberg 1995). Next, we performed a canonical analysis of principal components (CAP) ordination using Bray-Curtis distance for CLR-transformed bacterial and fungal communities. Genotype, Soil Inoculum, and Treatment were used to constrain the ordination and Sequencing Depth was partialled out to remove noise due to this technical nuisance variable. Then we used a linear model to determine whether the bacterial and fungal taxa counts varied by our explanatory variables.

## Results

### Heterosis for chlorophyll content and stem diameter in the agricultural and prairie inocula was affected in opposite directions

In Experiment 1, we grew plants with inocula derived from agricultural, prairie, or steam-sterilized (“control”) soil to determine if inoculum source had an effect on heterosis expression. We measured early plant traits, such as emergence, height, chlorophyll content, stem diameter, and biomass, over the course of four weeks then calculated heterosis for each trait. Here, we only report effects on better parent heterosis, but calculations for mid-parent heterosis are provided (Supplemental Table 1). After two weeks of growth, we did not observe a difference in the proportion of emerged plants due to inocula (Supplemental Figure 1). For week 2 chlorophyll content, we observed an increase in heterosis for the control (steam-sterilized) soil relative to the agriculture and prairie inocula (Supplemental Figure 2 and Figure 3), and for shoot biomass relative to the prairie inocula (Supplemental Figure 2). In contrast, soil sterilization did not affect heterosis of plant height, biomass, or stem diameter relative to the agriculture or prairie inocula (Supplemental Figure 2 and Figure 3). Inocula source (agriculture vs. prairie) did not have a large effect on heterosis expression with only stem diameter and week 4 chlorophyll content (Figure 1) differing between agriculture and prairie inocula in opposite directions. Stem diameter was greatest in the agricultural inocula, as indicated by the positive value for observed ΔBPH, whereas chlorophyll content was greatest in the prairie inocula, indicated by a negative observed ΔBPH. These differences were largely due to an increase in heterosis for these traits in the prairie inocula. When examining heterosis for individual inocula, we observed the weakest heterosis across traits in the first agriculture inoculum (avg. BPH = 0.06; MPH = 0.16) and the greatest heterosis was observed in the first prairie inoculum (avg. BPH = 0.17; MPH = 0.22) (Supplemental Table 1).

**Figure 1.**
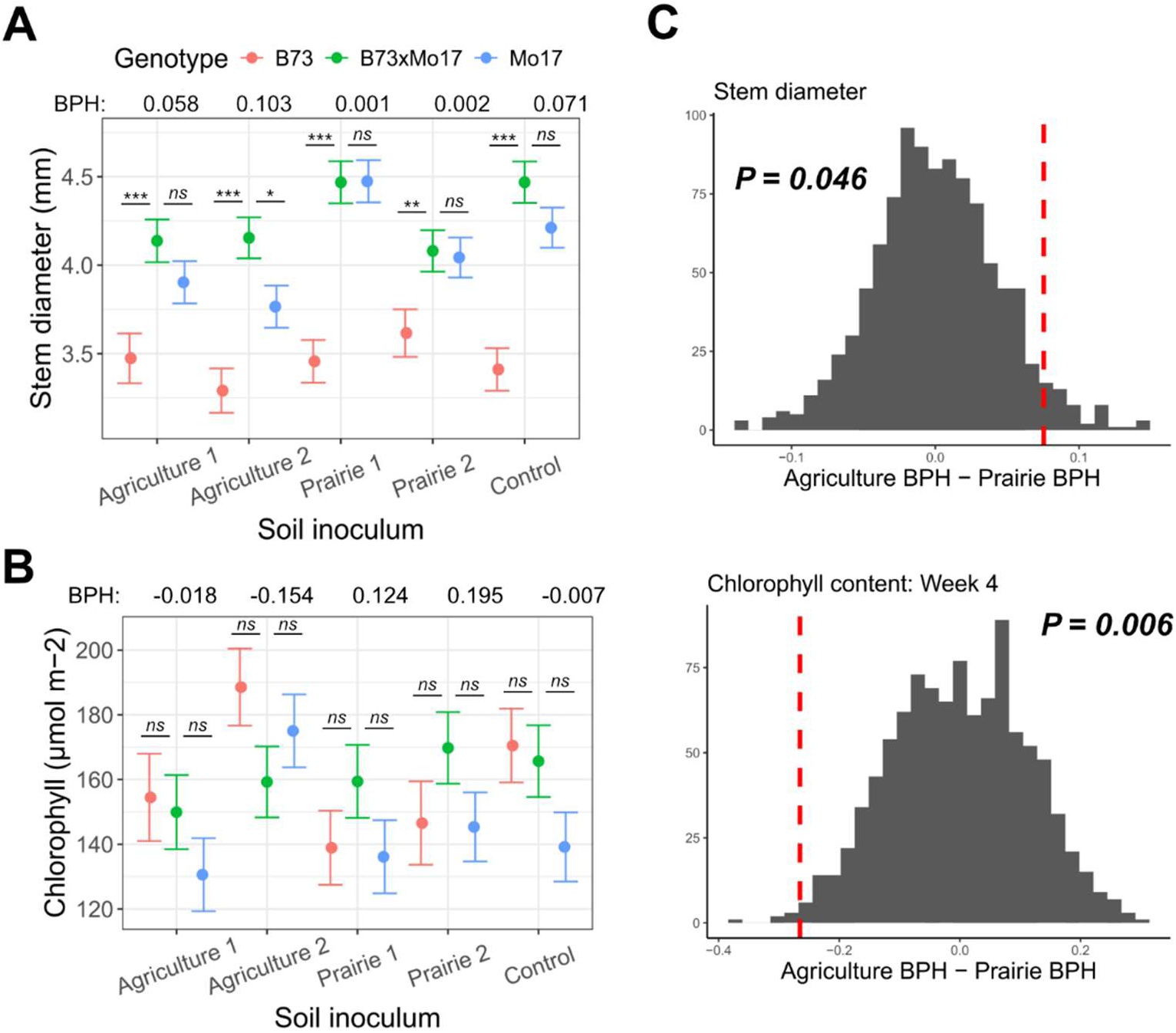
In Experiment 1, we grew maize with agriculture, prairie, or control soil under controlled growth chamber settings. After four weeks of growth, better parent heterosis (BPH) was greatest in the second agriculture soil for stem diameter (A) and greatest in the second prairie soil for week 4 chlorophyll content (B). Points show the estimated marginal mean (EMM) values for each genotype in each soil and error bars show the standard error for the EMMs. (C) BPH was calculated for soil inoculum using EMM values for stem diameter and week 4 chlorophyll content. The observed ΔBPH is shown as a vertical red line and the histogram shows the distributions of ΔBPH for 999 permutations of the data with respect to soil inocula.

### Low nutrient treatment results in strong heterosis across traits for agriculture inoculum

In Experiment 2, we employed a high and low nutrient treatment with the inoculum that displayed the weakest (agriculture 1) and greatest (prairie 1) average heterosis from Experiment 1. We measured early plant traits over the course of four weeks to determine if inocula and nutrient treatment have an effect on better parent heterosis expression. Similar to Experiment 1, we did not observe a difference in emergence proportion due to inocula or nutrient treatment (Supplemental Figure 5). For plant traits other than emergence, we observed increased trait values under the high nutrient treatment regardless of plant genotype (Figure 2). However, we also observed an effect of nutrient treatment on heterosis for stem diameter for both inocula (Supplemental Figures 6 and 7). Interestingly, heterosis for week two plant height (Figure 2) and chlorophyll content (Supplemental Figure 6) was affected by the nutrient treatment for only the agriculture inoculum in opposite directions. In general, heterosis was more affected by the nutrient treatment in combination with the agricultural inoculum (Supplemental Figure 6). With the exception of week two chlorophyll content, this was driven by a decrease in heterosis under the high nutrient relative to the low nutrient treatment, suggesting an interaction between agricultural soil microorganisms and nutrient amendment.

**Figure 2.**
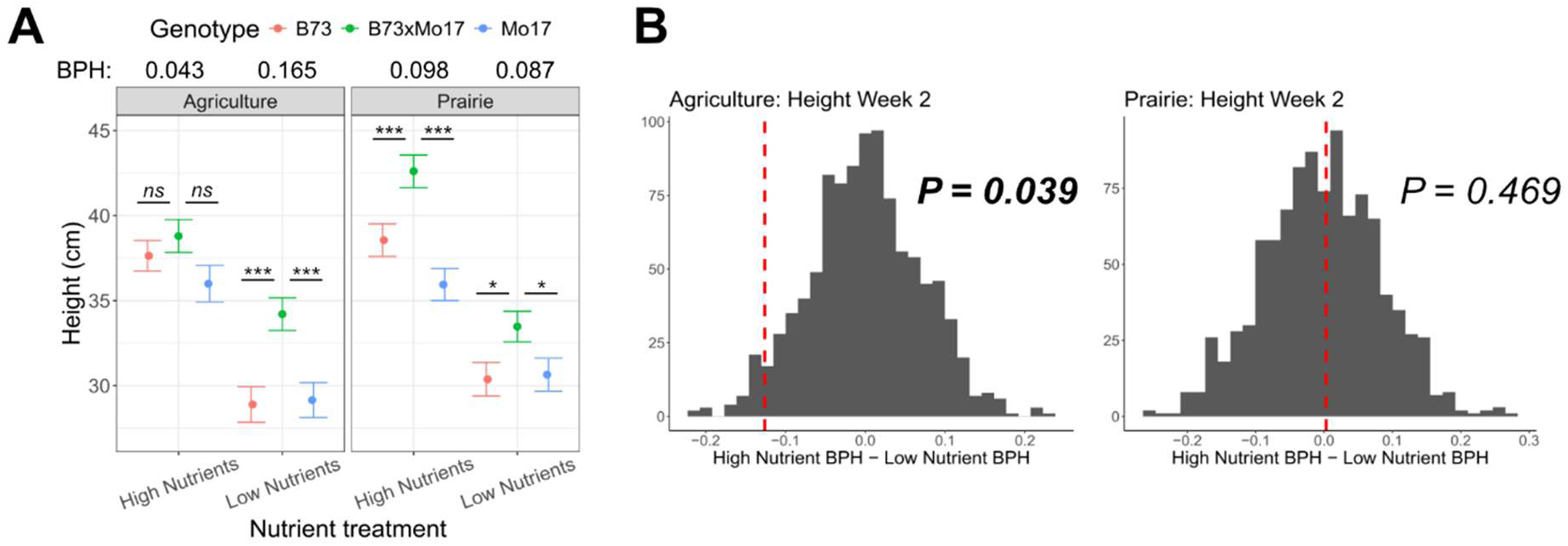
In Experiment 2, we grew maize with agriculture or prairie soil and a high or low nutrient treatment. After two weeks of growth, betterparent heterosis (BPH) of (A) plant height was greatest in the low nutrient treatment for the agriculture soil but greatest in the high nutrient treatment for the prairie soil. Error bars show the standard error for the estimated marginal mean values. (B) BPH was calculated for each soil using the EMM trait values. The observed ΔBPH is shown as a vertical red line and the histogram shows the distributions of ΔBPH for 999 permutations of the data with respect to nutrient treatment.

### Soil pH and total percent leaf nitrogen is higher for inbred genotypes

In Experiment 2, we also measured soil pH at two and four weeks to examine the effect of nutrient addition on soil chemistry. We did not observe an effect after two weeks but after four weeks, we observed lower soil pH across genotypes in the high nutrient relative to the low nutrient treatments (p=<0.01, F_1,140_=33). This suggests that the nutrient treatment was directly responsible for altering the soil pH over the course of the experiment. Interestingly, we also observed an effect of genotype after four weeks (p=<0.05, F_2,137_=5), in which soil was less acidic in pots with the inbred genotypes than pots with the hybrid, regardless of soil inocula (Figure 3). We also observed negative heterosis for leaf nitrogen content, in which the inbred genotypes contained a greater nitrogen concentration in the leaf tissue than the hybrid although not significantly different (Figure 3). Total percent carbon in leaf tissue was also strongly affected by plant genotype (p=<0.01, F_2,9_=10), as well as nutrient treatment (p=<0.01, F_1,6_=9), but a clear heterotic pattern was not apparent.

**Figure 3.**
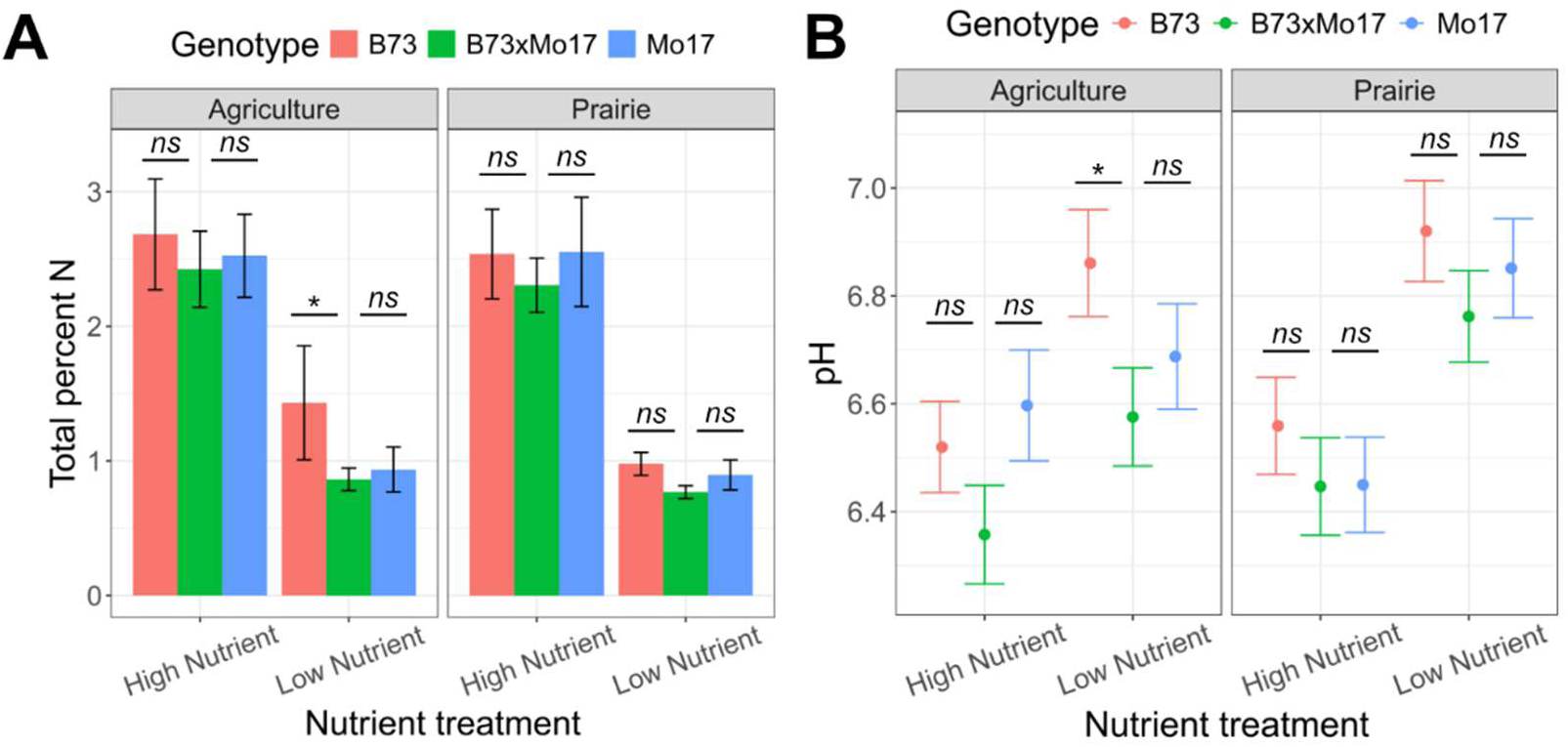
(A) Total percent nitrogen in shoot tissue was strongly affected by nutrient treatment (P=<0.001) with a greater percent nitrogen across genotypes in the high nutrient treatment. Error bars represent standard deviation. (B) Soil pH was also strongly affected by nutrient treatment after four weeks (P=<0.001), in which pH was decreased in the high nutrient treatment. Error bars show the standard error for the estimated marginal mean values.

### Soil inocula and nutrient treatment drive bacterial and fungal community composition

To assess the effects of soil inocula and nutrient treatment, we characterized the bacterial and fungal community of the root endosphere. The diversity of bacterial ASVs was significantly affected by nutrient treatment as determined by the Shannon Index and Inverse Simpson (Supplemental Table 2). In addition, bacterial diversity was significantly affected by genotype for the Shannon Index but genotype only marginally explained bacterial diversity according to Inverse Simpson. In contrast, diversity of fungal ASVs was not significantly affected by any of our explanatory variables (Supplemental Table 3). A canonical analysis of principal coordinates (CAP) ordination constrained by Genotype, Soil Inoculum, and Treatment revealed that bacteria community composition was significantly affected by Soil Inoculum and Treatment, as well as their interaction (Figure 4). We observed a similar trend for fungal community composition, in that Soil Inoculum and Treatment were significant, but their interaction was only marginally significant.

**Figure 4.**
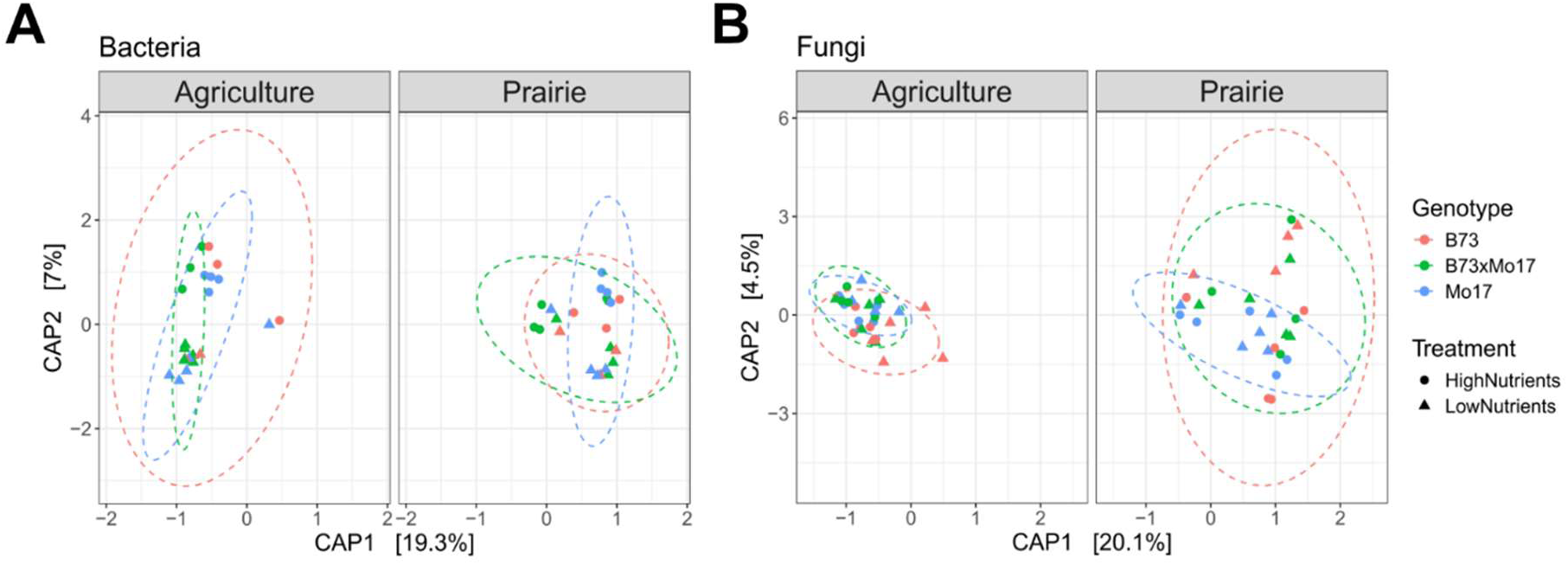
CAP ordination for bacterial (A) and fungal (B) root endosphere communities constrained by soil inocula, genotype, and nutrient treatment. Bacterial community composition was strongly driven by soil inocula and nutrient treatment (P<0.001) and their interaction (P<0.05). Fungal community composition was also driven by soil inocula (P<0.001) and nutrient treatment (P<0.05).

## Discussion

In Experiment 1, we observed similar results reported in previous work (Wagner et al. 2021), in which soil sterilization increased better parent heterosis for root biomass relative to the Kansas agricultural soil (Supplemental Table 1). In fact, the weakest heterosis across traits was observed in the first agricultural soil, which was collected from the same site where the previous Kansas field experiment was conducted (Wagner et al. 2021). Here the control soil resulted in an increase in heterosis, while the first agricultural soil inoculum exhibited a decrease in heterosis. This increase in heterosis appears to be due to a decrease in performance of the inbreds, rather than an increase in performance of the hybrid. In contrast, the second prairie inoculum exhibited greater heterosis across traits than the control soil. This reaffirms the observation that microbial effects on heterosis are likely specific to the local microbial community (Wagner et al. 2021). We hypothesized that pathogen buildup in the agricultural soil would result in a decrease in performance of the inbreds and therefore, an increase in better parent heterosis expression. However, both the agricultural and prairie inocula exhibited similar heterosis expression across traits. In total, only two (stem diameter and week 4 chlorophyll content) out of seven traits exhibited contrasting heterosis between the two inoculum types. These differences were not consistent, however, since stem diameter heterosis was greatest in the agricultural inocula while chlorophyll content heterosis was greatest in the prairie inocula. Assuming that the agricultural soils were indeed richer in pathogens, these results contradict one of the proposed mechanisms for microbe-dependent heterosis, which suggests that hybrids may be more resistant to pathogenic soil microbes than inbreds. Therefore, there may be other environmental factors contributing to variation in heterosis expression rather than soil pathogens.

In Experiment 2, we employed a high and low nutrient treatment to explore the interaction between nutrient status and soil inoculum on heterosis expression. Since the agricultural soil location has a legacy of fertilization, we expected this interaction to be stronger in the agricultural inoculum. Unsurprisingly, we observed an increase in all plant traits under the high nutrient treatment regardless of plant genotype or soil inoculum. Interestingly, all traits besides week 4 chlorophyll content were more affected by the nutrient treatment in the agricultural than in the prairie inoculum. In total, three out of eight traits were significantly affected by the nutrient treatment in the agricultural inoculum, which was due to a decrease in heterosis under the high nutrient treatment. In addition, the interaction between genotype, soil, and nutrient treatment were significant for two (height week 2 and chlorophyll content week 4) of these traits and marginally significant for the third (stem diameter week 4). Due to these results and the fact that the inocula contributed only 15% v/v of each total soil, it can be suggested that this effect was driven by the interaction between agricultural soil microorganisms and the nutrient amendment rather than agricultural chemical soil properties.

Nutrient availability, especially nitrogen, is a strong predictor of global soil microbial community structure (Wang et al., 2023). Further, the addition of nutrients can affect the potential benefit of microbial symbiosis for plants. For example, mycorrhizal fungi increase the shoot biomass of maize under phosphorus-limited conditions, but decrease biomass when phosphorus is plentiful (Kaeppler et al., 2000). Since the agricultural soil location has a legacy of fertilization, we expected this interaction to be stronger in the agricultural inoculum. We observed an effect of soil and nutrient treatment on the composition of both the bacterial and fungal communities in the root endosphere. In addition, the interaction between soil and treatment significantly affected bacterial community composition but not the fungal community. However, alpha diversity of the bacterial and fungal community was only affected by nutrient treatment. Since the nutrient treatment strongly altered soil pH, which is a strong indicator of microbiome composition (Lauber et al. 2009), it is unsurprising that treatment affected both diversity and composition of the bacteria and fungal community. In contrast, we only observed a marginal effect of genotype on root bacterial composition and Inverse Simpson diversity, but a significant effect for Shannon diversity. However, the root fungal composition and diversity were not affected by genotype at all.

Interestingly, we also observed an effect of genotype on soil pH after four weeks of growth, in which pH was more acidic in pots with hybrid genotypes relative to inbred genotypes. During active growth, plants increase acidity along their roots since cation uptake exceeds anion uptake (Xu et al. 2006). Because the hybrids exhibited greater growth across traits, it can be inferred that this resulted in more acidic soil pH for pots with hybrid genotypes. Consequently, this could also contribute to the differences in bacterial diversity we observed due to genotype since bacteria are highly sensitive to pH. We also observed an increase in leaf nitrogen content strongly affected by nutrient treatment, in which leaf nitrogen content was increased in the high nutrient treatment regardless of plant genotype. We did not observe a significant effect of genotype on leaf nitrogen content, however, the inbred genotypes appear to have slightly more nitrogen accumulation than the hybrid regardless of soil inoculum or nutrient treatment. Total percent carbon in leaf tissue was strongly affected by plant genotype, as well as nutrient treatment, but a clear effect on heterosis was not apparent.

Together, our results suggest that the agriculture and prairie soil inocula we use here do not differ in their effect on heterosis and do not affect heterosis in a consistent way. However, these conclusions are based on the analysis of just two soils tested per inoculum type. Since the soils were collected from northeast Kansas, regional climatic conditions could also result in a diluting effect of microbial community dissimilarity between the two inocula types. For fungi, climatic factors are the best global predictors of richness and community composition (Tedersoo et al., 2014) and therefore, may not substantially differ between our soils.

In contrast, the introduction of nutrients, which are historically common for the agricultural soil, resulted in a larger effect on heterosis expression. Specifically, we observed an effect of nutrient treatment on heterosis for the agricultural inoculum, in which heterosis was decreased in the high nutrient relative to low nutrient treatment. This effect on heterosis was driven by greater variation in hybrid performance due to soil inocula in the high nutrient treatment (Supplemental Figure 8); in contrast, the hybrid was less responsive to the soil inocula than the inbreds in the low nutrient treatment. We also observed an effect of soil inoculum, nutrient treatment, and their interaction on bacterial community composition and diversity. In addition, genotype and the interaction between soil and genotype was marginally significant for bacteria community composition. Together, this suggests that the nutrient amendment simultaneously had an effect on the root microbiome and heterosis expression.

These results further demonstrate that the soil microbial community may play a role in heterosis expression but soil nutrients are likely the stronger driver. They also provide direct evidence for the interactive effect of the soil microbial community and abiotic environment on heterosis expression, which was previously undescribed. Future work will require experimentation to determine the effects of this interaction on the molecular and physiological mechanisms of heterosis. This research may lead to the development of new or advanced applications of microbiome sciences, including the genetic improvement of plant phenotypic response to the soil microbiomes (Clouse and Wagner, 2021). Ultimately, a great deal of research is needed to advance our understanding of heterosis across crop species and how it may improve agricultural sustainability generally.

## Acknowledgements

We thank Aoesta Rudick for generating the C:N data. This work was supported by the National Science Foundation grant #IOS-2033621 to MRW, PBK, and MK. KMC was also supported by a fellowship from the National Institute for Food and Agriculture, grant #2023-67011-40397. We thank the University of Kansas Medical Center Genomics Core for generating the sequence data sets. The Genomics Core is supported by the Kansas Intellectual and Developmental Disabilities Research Center (NIH U54 HD 090216), the Molecular Regulation of Cell Development and Differentiation – COBRE (P30 GM122731), the NIH S10 High End Instrumentation Grant (NIH S10OD021743) and the Frontiers CTSA Grant (UL1TR002366).

## Supplemental Information

**Supplemental Figure 1.**
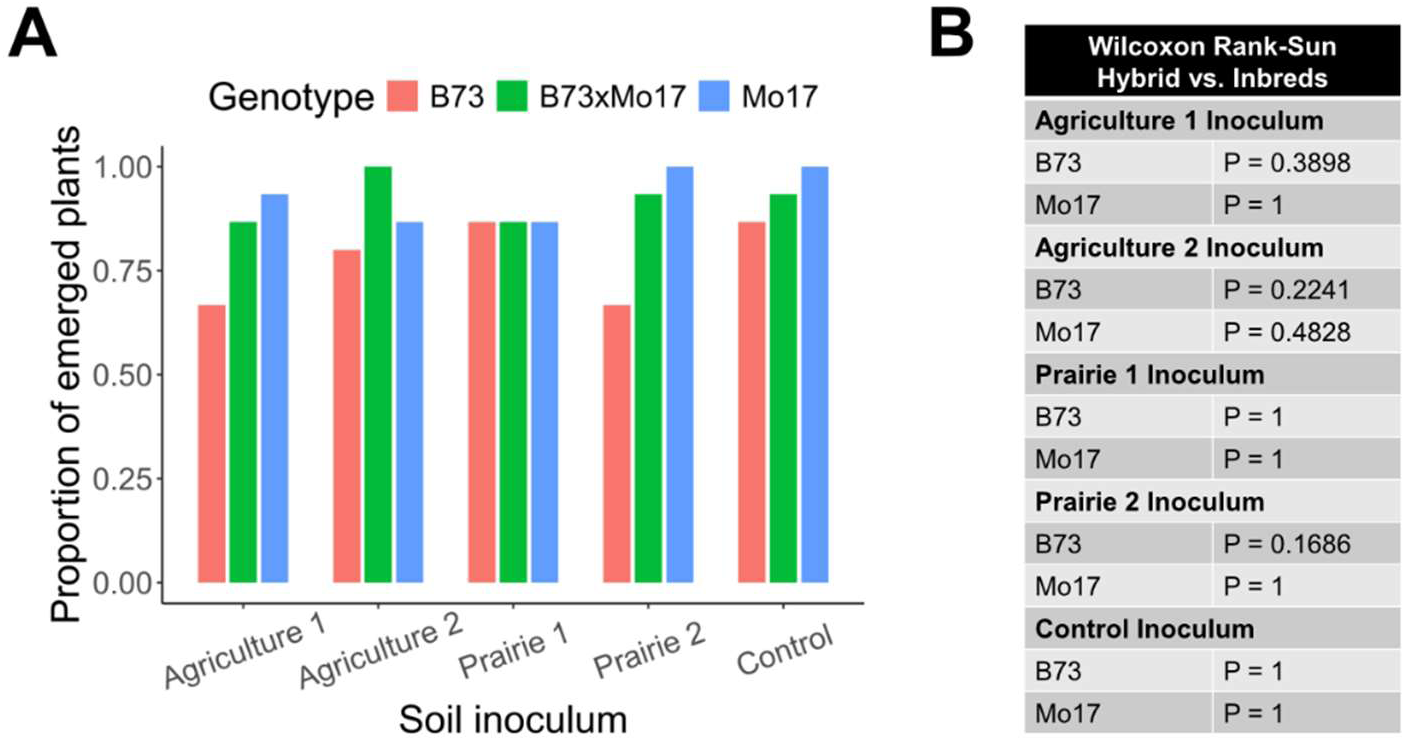
(A) Mean emergence proportions for each genotype after 14-days in each soil inoculum. (B) Wilcoxon rank-sum tests for pairwise comparisons of each inbred to the hybrid for each soil inoculum. P-values were adjusted to correct for multiple comparisons using the Benjamini-Hochberg method.

**Supplemental Figure 2.**
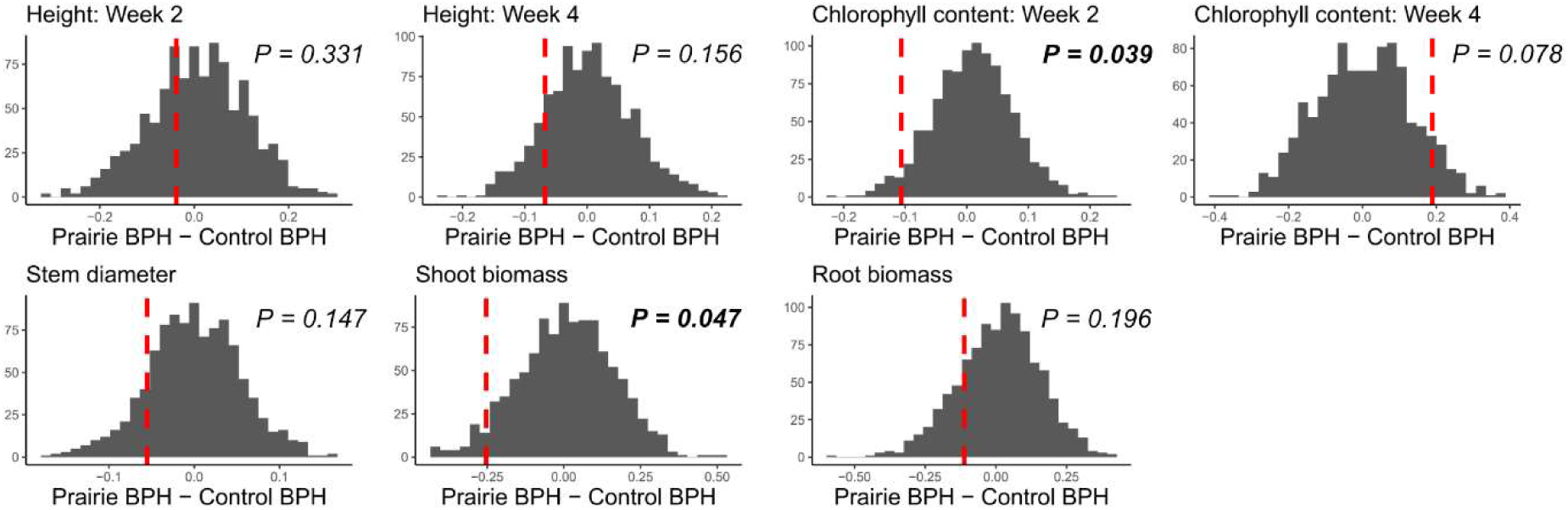
Better-parent heterosis (BPH) was calculated for prairie versus control soil inoculum using EMM values for each trait. The observed ΔBPH is shown as a vertical red line and the histogram shows the distributions of ΔBPH for 999 permutations of the data with respect to soil.

**Supplemental Figure 3.**
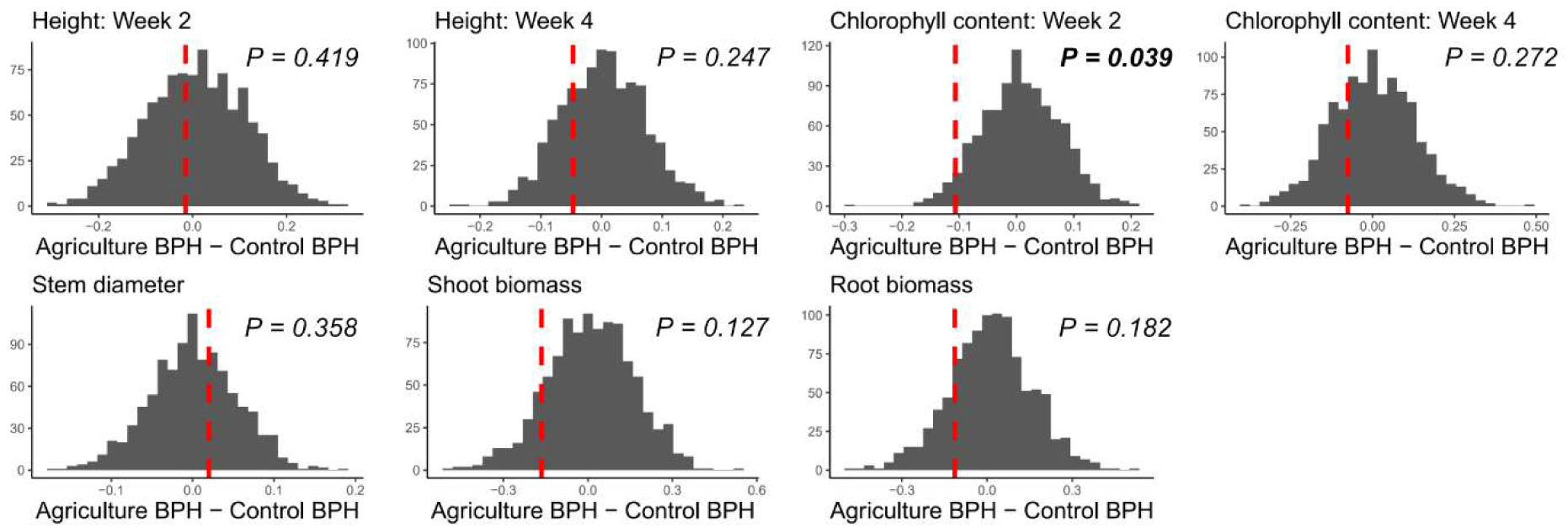
Better-parent heterosis (BPH) was calculated for agriculture versus control soil inoculum using EMM values for each trait. The observed ΔBPH is shown as a vertical red tine and the histogram shows the distributions of ΔBPH for 999 permutations of the data with respect to soil.

**Supplemental Figure 4.**
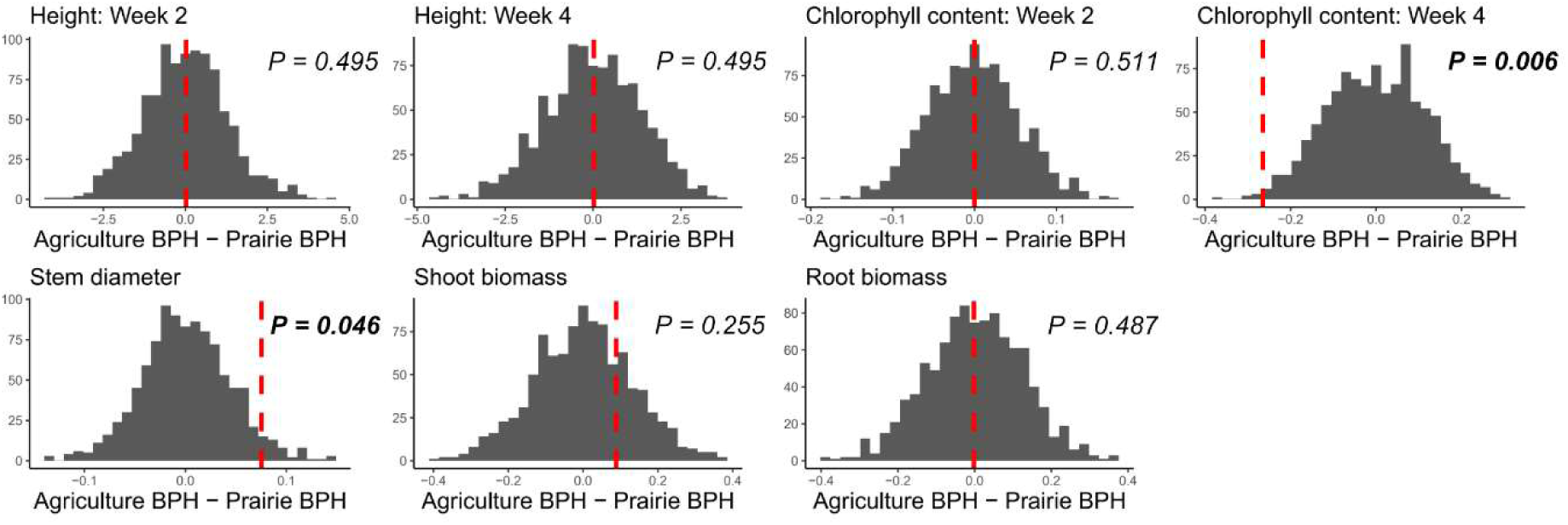
Better-parent heterosis (BPH) was calculated for agriculture versus prairie soil inoculum using EMM values for each trail. The observed ΔBPH is shown as a vertical red line and the histogram shows the distributions of ΔBPH for 999 permutations of the data with respect to soil.

**Supplemental Figure 5.**
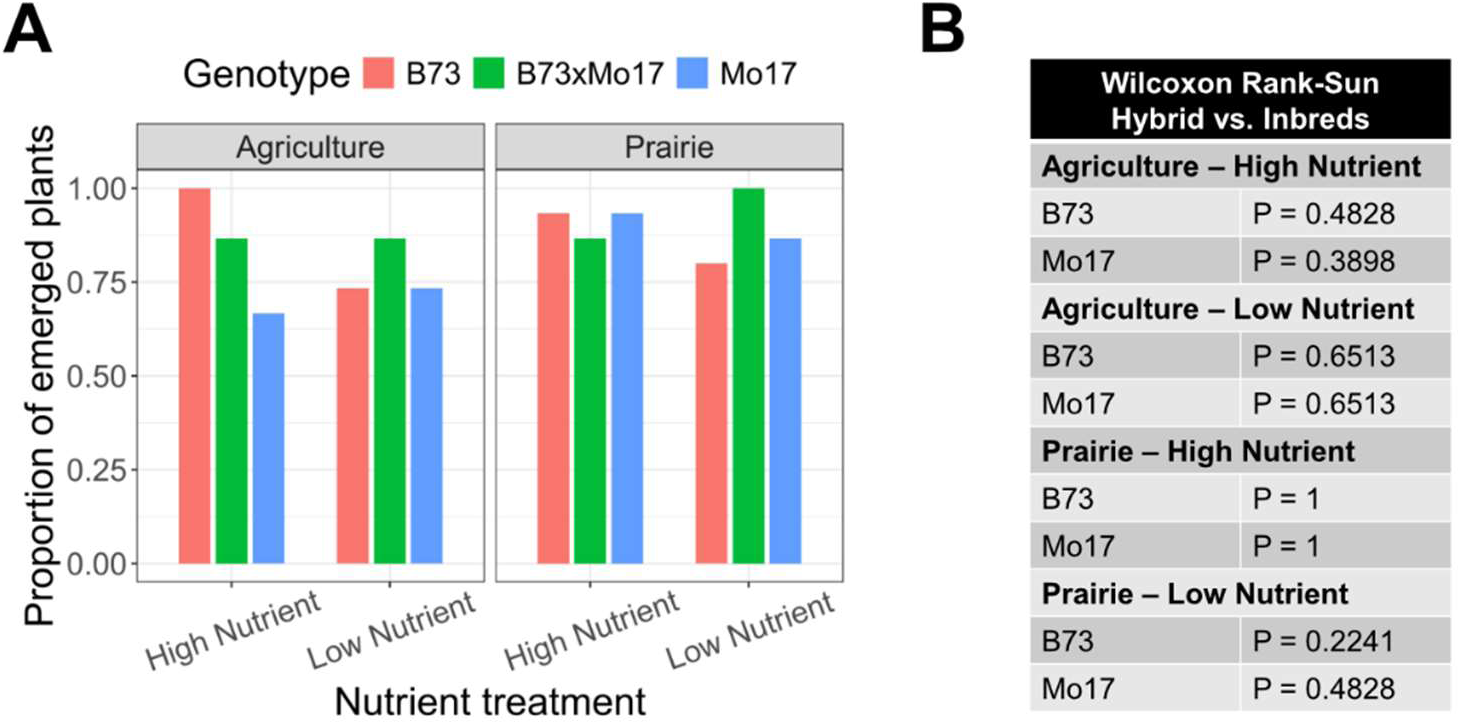
(A) Mean emergence proportions for each genotype after 14-days in each soil and nutrient treatment combination. (B) Wilcoxon rank-sum tests for pairwise comparisons of each inbred to the hybrid for each soil and nutrient treatment combination. P-values were adjusted to correct for multiple comparisons using the Benjamini-Hochberg method.

**Supplemental Figure 6.**
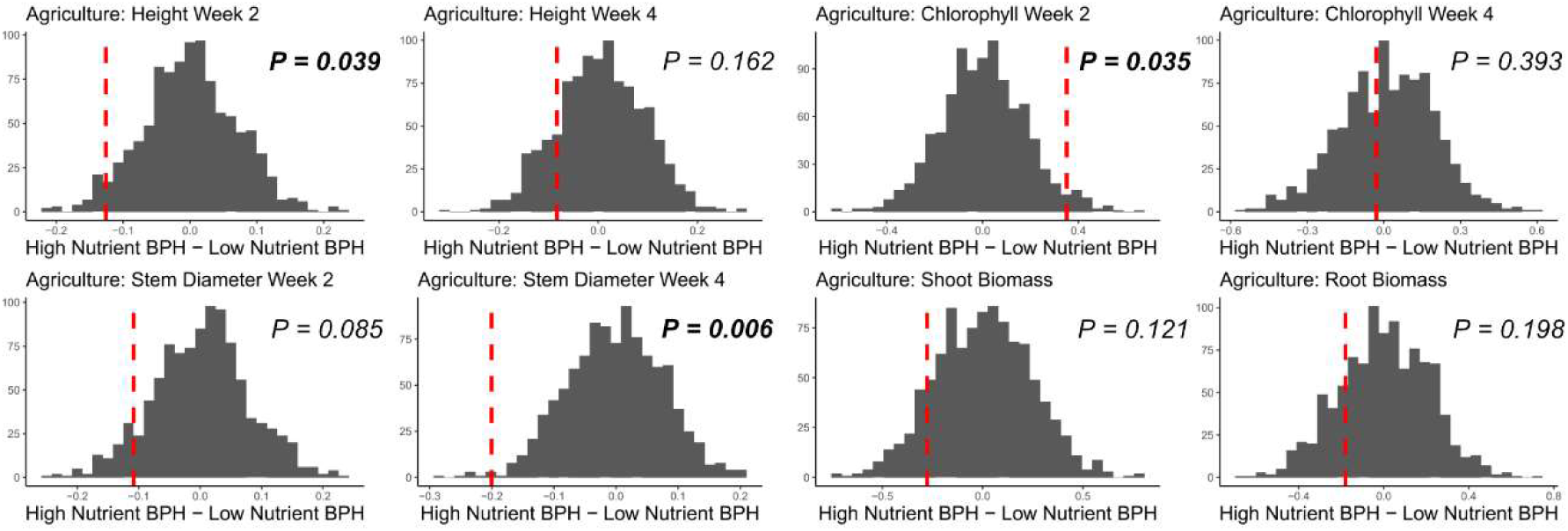
Better-parent heterosis (BPH) was calculated for high versus low nutrient treatment for the agricultual inoculum using EMM values for each trait. The observed ΔBPH is shown as a vertical red line and the histogram shows the distributions of ΔBPH for 999 permutations of the data with respect to nutrient treatment.

**Supplemental Figure 7.**
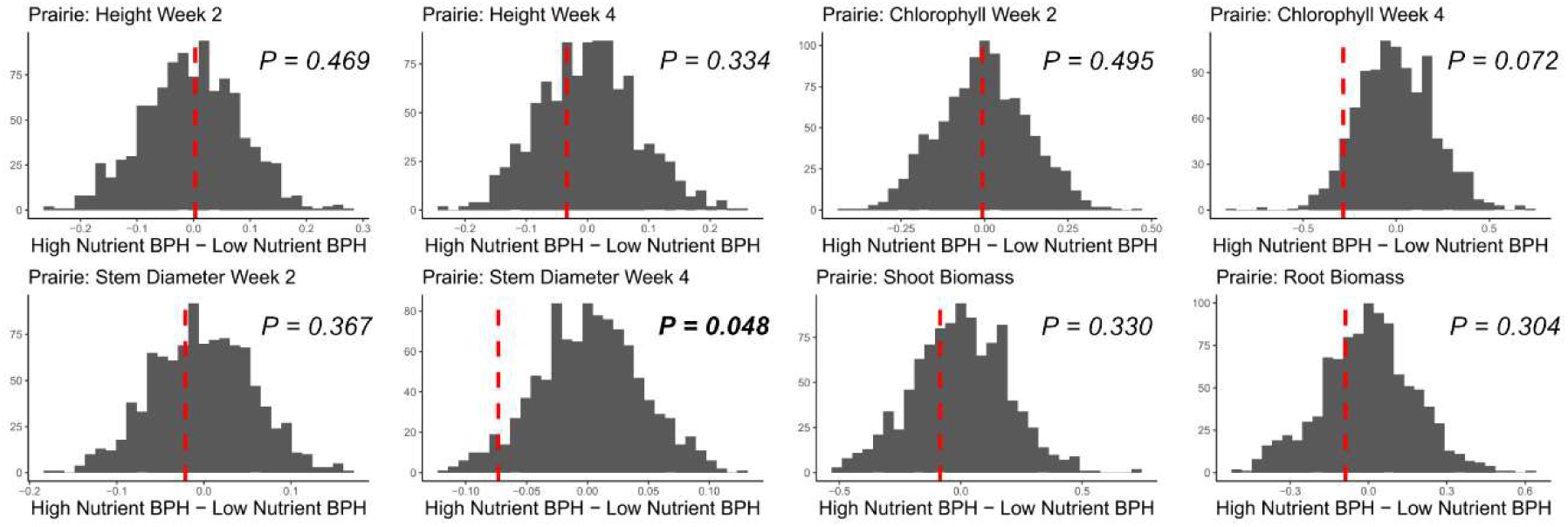
Better-parent heterosis (BPH) was calculated for high versus low nutrient treatment for the prairie inoculum using EMM values for each trait. The observed ΔBPH is shown as a vertical red line and the histogram shows the distributions of ΔBPH for 999 permutations of the data with respect to nutrient treatment.

**Supplemental Figure 8.**
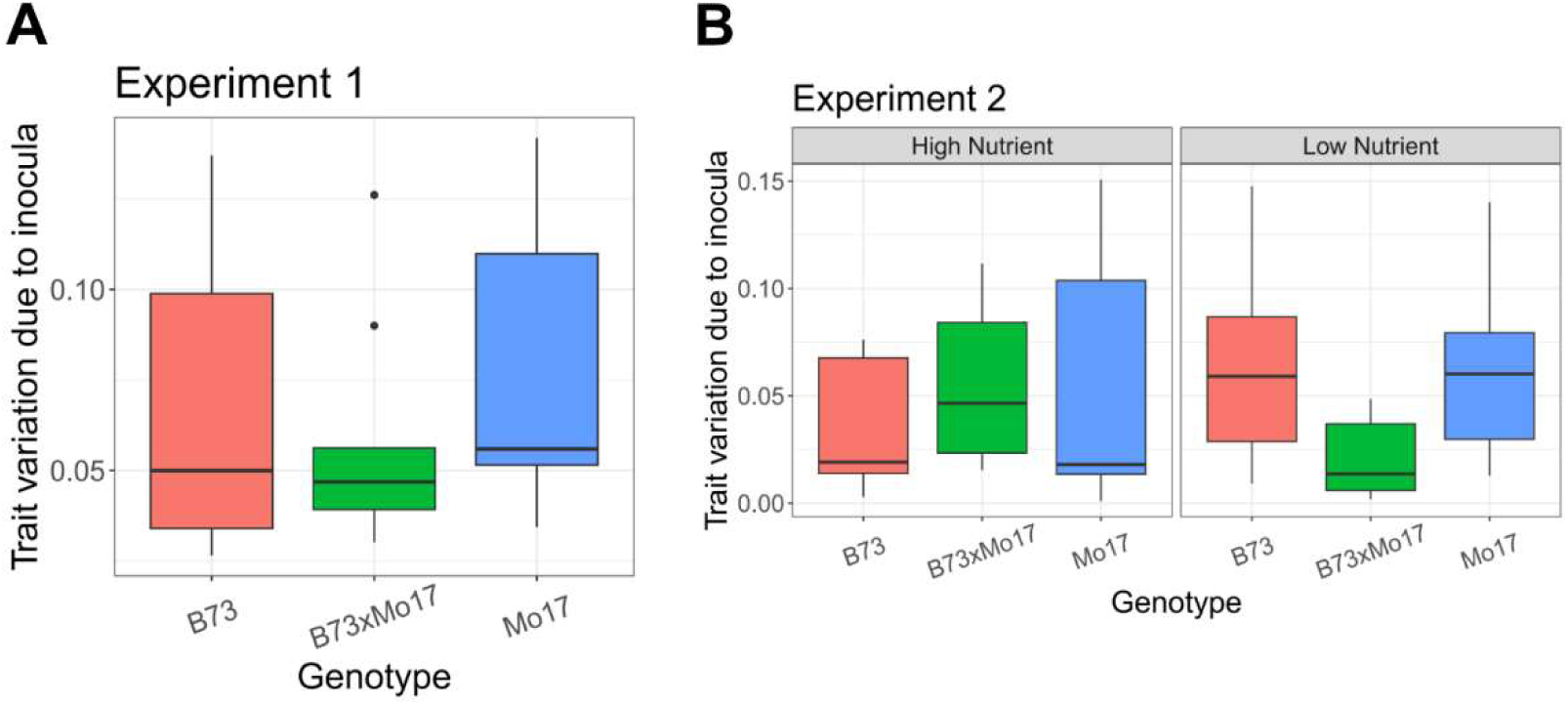
To determine the amount of variation in each genotype’s phenotypic response due to soil inocula, we calculated the coefficient of variation across plant traits using their estimated marginal mean value. The coefficient of variation for each genotype across plant traits (9 total) and soil inocula in Experiment 1 (A) and for each genotype and nutrient treatment across plant traits (8 total) and soil inocula in Experiment 2 (B)

**Supplemental Figure 9.**
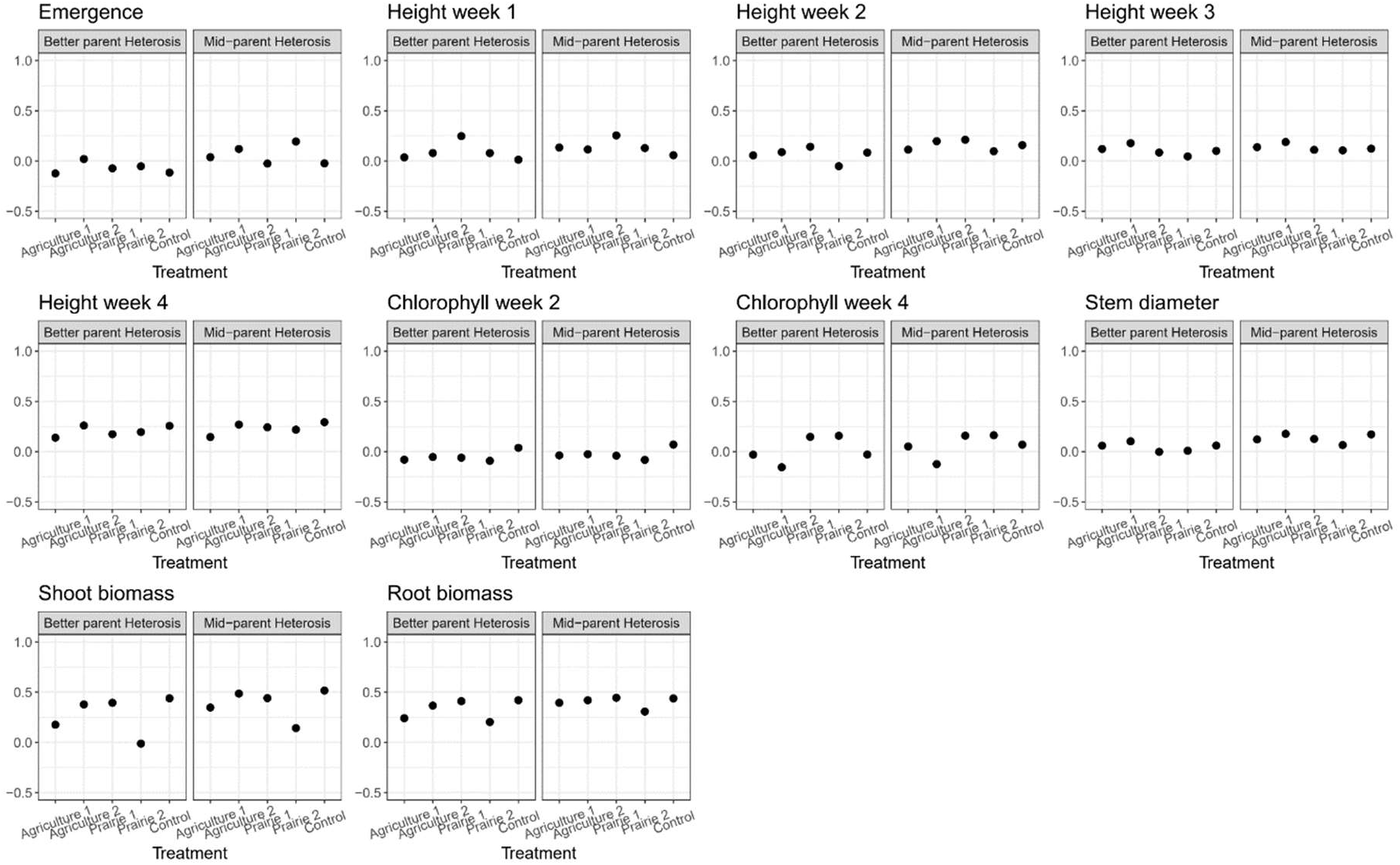
Better parent and mid-parent calculations for each plant trait in each soil inoculum for Experiment 1.

**Supplemental Figure 10.**
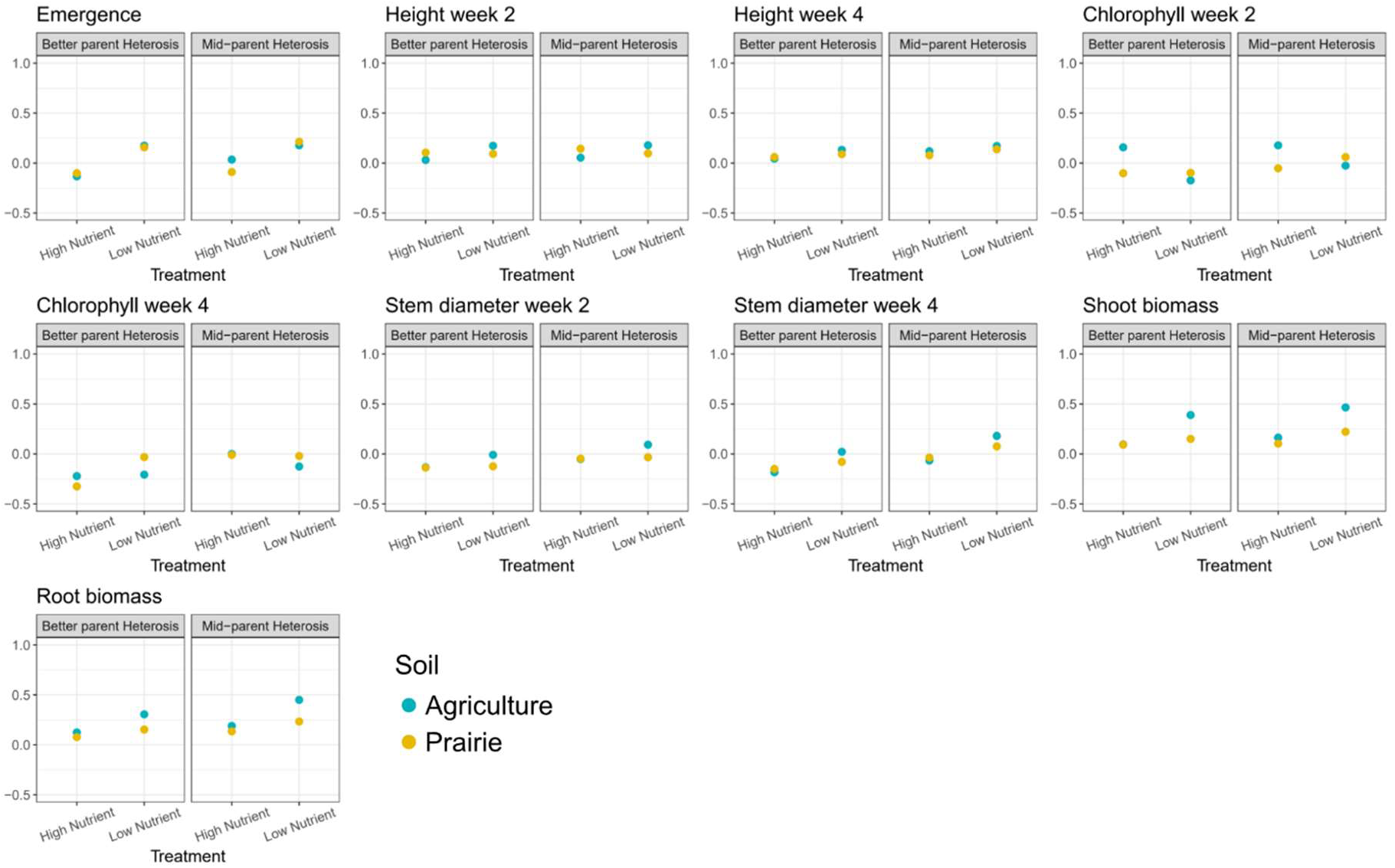
Better parent and mid-parent calculations for each plant trait in each soil inoculum and nutrient treatment for Experiment 2.

**Supplemental Table 1.** Betterparent and mid-parent heterosis calculations for each plant trait in each soil inoculum for Experiment 1.

| Soil Inoculum | Plant Trait | Betterparent Heterosis | Mid-parent Heterosis |
| --- | --- | --- | --- |
| Agriculture 1 | Emergence | -0.167 | 0.000 |
| Agriculture 1 | HeightWeek1 | 0.028 | 0.124 |
| Agriculture 1 | HeightWeek2 | 0.083 | 0.142 |
| Agriculture 1 | HeightWeek3 | 0.123 | 0.135 |
| Agriculture 1 | HeightWeek4 | 0.085 | 0.106 |
| Agriculture 1 | ChlorophyllWeek2 | -0.080 | -0.037 |
| Agriculture 1 | ChlorophyllWeek4 | -0.018 | 0.076 |
| Agriculture 1 | StemDiameter | 0.058 | 0.121 |
| Agriculture 1 | ShootBiomass | 0.140 | 0.340 |
| Agriculture 1 | RootBiomass | 0.208 | 0.415 |
| Agriculture 2 | Emergence | -0.087 | 0.050 |
| Agriculture 2 | HeightWeek1 | 0.088 | 0.123 |
| Agriculture 2 | HeightWeek2 | 0.120 | 0.228 |
| Agriculture 2 | HeightWeek3 | 0.193 | 0.206 |
| Agriculture 2 | HeightWeek4 | 0.298 | 0.305 |
| Agriculture 2 | ChlorophyllWeek2 | -0.053 | -0.025 |
| Agriculture 2 | ChlorophyllWeek4 | -0.154 | -0.124 |
| Agriculture 2 | StemDiameter | 0.103 | 0.170 |
| Agriculture 2 | ShootBiomass | 0.371 | 0.494 |
| Agriculture 2 | RootBiomass | 0.344 | 0.426 |
| Control | Emergence | -0.154 | -0.043 |
| Control | HeightWeek1 | 0.028 | 0.065 |
| Control | HeightWeek2 | 0.096 | 0.173 |
| Control | HeightWeek3 | 0.121 | 0.142 |
| Control | HeightWeek4 | 0.282 | 0.330 |
| Control | ChlorophyllWeek2 | 0.039 | 0.071 |
| Control | ChlorophyllWeek4 | -0.007 | 0.086 |
| Control | StemDiameter | 0.071 | 0.176 |
| Control | ShootBiomass | 0.442 | 0.546 |
| Control | RootBiomass | 0.466 | 0.515 |
| Prairie 1 | Emergence | -0.130 | -0.048 |
| Prairie 1 | HeightWeek1 | 0.261 | 0.262 |
| Prairie 1 | HeightWeek2 | 0.175 | 0.236 |
| Prairie 1 | HeightWeek3 | 0.087 | 0.128 |
| Prairie 1 | HeightWeek4 | 0.179 | 0.274 |
| Prairie 1 | ChlorophyllWeek2 | -0.060 | -0.040 |
| Prairie 1 | ChlorophyllWeek4 | 0.124 | 0.138 |
| Prairie 1 | StemDiameter | 0.001 | 0.137 |
| Prairie 1 | ShootBiomass | 0.400 | 0.432 |
| Prairie 1 | RootBiomass | 0.403 | 0.417 |
| Prairie 2 | Emergence | -0.040 | 0.263 |
| Prairie 2 | HeightWeek1 | 0.069 | 0.126 |
| Prairie 2 | HeightWeek2 | -0.062 | 0.096 |
| Prairie 2 | HeightWeek3 | 0.051 | 0.107 |
| Prairie 2 | HeightWeek4 | 0.183 | 0.190 |
| Prairie 2 | ChlorophyllWeek2 | -0.091 | -0.082 |
| Prairie 2 | ChlorophyllWeek4 | 0.195 | 0.197 |
| Prairie 2 | StemDiameter | 0.002 | 0.059 |
| Prairie 2 | ShootBiomass | 0.037 | 0.176 |
| Prairie 2 | RootBiomass | 0.283 | 0.348 |

**Supplemental Table 2.** Shannon and Inverse Simpson alpha diversity indices for bacteria. P-values were adjusted to correct for multiple comparisons using the Benjamini-Hochberg method.

| Response | Term | Numerator Degrees of Freedom | Denominator Degrees of Freedom | F | P-adjusted |
| --- | --- | --- | --- | --- | --- |
| Shannon | Soil | 1 | 32 | 0.00 | 0.973 |
| Shannon | Genotype | 2 | 32 | 2.06 | 0.207 |
| Shannon | Treatment | 1 | 32 | 16.17 | 0.001 |
| Shannon | Sequencing Depth | 1 | 32 | 18.62 | 0.000 |
| Shannon | Soil:Genotype | 2 | 32 | 1.58 | 0.222 |
| Shannon | Soil:Treatment | 1 | 32 | 1.49 | 0.231 |
| Shannon | Genotype: Treatment | 2 | 32 | 1.14 | 0.665 |
| Shannon | Soil:Genotype:Treatment | 2 | 32 | 0.80 | 0.467 |
| InvSimpson | Soil | 1 | 32 | 1.27 | 0.535 |
| InvSimpson | Genotype | 2 | 32 | 1.65 | 0.207 |
| InvSimpson | Treatment | 1 | 32 | 7.64 | 0.009 |
| InvSimpson | Sequencing Depth | 1 | 32 | 5.60 | 0.024 |
| InvSimpson | Soil:Genotype | 2 | 32 | 2.18 | 0.222 |
| InvSimpson | Soil:Treatment | 1 | 32 | 2.89 | 0.197 |
| InvSimpson | Genotype:Treatment | 2 | 32 | 0.22 | 0.803 |
| InvSimpson | Soil:Genotype:Treatment | 2 | 32 | 0.78 | 0.467 |

**Supplemental Table 3.** Shannon and Inverse Simpson alpha diversity indices for fungi. P-values were adjusted to correct for multiple comparisons using the Benjamini-Hochberg method.

| Response | Term | Numerator Degrees of Freedom | Denominator Degrees of Freedom | F | P-adjusted |
| --- | --- | --- | --- | --- | --- |
| Shannon | Soil | 1 | 43.29 | 0.08 | 0.784 |
| Shannon | Genotype | 2 | 44.25 | 0.99 | 0.675 |
| Shannon | Treatment | 1 | 44.39 | 0.13 | 0.718 |
| Shannon | Sequencing Depth | 1 | 44.99 | 14.29 | 0.000 |
| Shannon | Soil:Genotype | 2 | 44.76 | 0.07 | 0.933 |
| Shannon | Soil:Treatment | 1 | 44.99 | 0.49 | 0.688 |
| Shannon | Genotype: Treatment | 2 | 44.85 | 1.12 | 0.335 |
| Shannon | Soil:Genotype:Treatment | 2 | 44.00 | 0.77 | 0.491 |
| InvSimpson | Soil | 1 | 45.00 | 0.41 | 0.784 |
| InvSimpson | Genotype | 2 | 45.00 | 0.40 | 0.675 |
| InvSimpson | Treatment | 1 | 45.00 | 0.28 | 0.718 |
| InvSimpson | Sequencing Depth | 1 | 45.00 | 21.03 | 0.000 |
| InvSimpson | Soil:Genotype | 2 | 45.00 | 0.27 | 0.933 |
| InvSimpson | Soil:Treatment | 1 | 45.00 | 0.16 | 0.688 |
| InvSimpson | Genotype:Treatment | 2 | 45.00 | 1.73 | 0.335 |
| InvSimpson | Soil:Genotype:Treatment | 2 | 45.00 | 0.72 | 0.491 |

**Supplemental Table 4.** Betterparent and mid-parent heterosis calculations for each plant trait in each soil inoculum and nutrient treatment for Experiment 2.

| Soil Inoculum | Nutrient Treatment | Plant Trait | Betterparent Heterosis | Mid-parent Heterosis |
| --- | --- | --- | --- | --- |
| Agriculture | High Nutrient | ChlorophyllWeek2 | 0.162 | 0.176 |
| Agriculture | High Nutrient | ChlorophyllWeek4 | -0.222 | -0.003 |
| Agriculture | High Nutrient | Emergence | -0.133 | 0.040 |
| Agriculture | High Nutrient | HeightWeek2 | 0.043 | 0.069 |
| Agriculture | High Nutrient | HeightWeek4 | 0.053 | 0.127 |
| Agriculture | High Nutrient | RootBiomass | 0.124 | 0.186 |
| Agriculture | High Nutrient | ShootBiomass | 0.104 | 0.172 |
| Agriculture | High Nutrient | StemDiameterWeek2 | -0.123 | -0.038 |
| Agriculture | High Nutrient | StemDiameterWeek4 | -0.174 | -0.054 |
| Agriculture | Low Nutrient | ChlorophyllWeek2 | -0.193 | -0.039 |
| Agriculture | Low Nutrient | ChlorophyllWeek4 | -0.190 | -0.108 |
| Agriculture | Low Nutrient | Emergence | 0.182 | 0.182 |
| Agriculture | Low Nutrient | HeightWeek2 | 0.165 | 0.168 |
| Agriculture | Low Nutrient | HeightWeek4 | 0.133 | 0.167 |
| Agriculture | Low Nutrient | RootBiomass | 0.305 | 0.457 |
| Agriculture | Low Nutrient | ShootBiomass | 0.392 | 0.474 |
| Agriculture | Low Nutrient | StemDiameterWeek2 | -0.020 | 0.085 |
| Agriculture | Low Nutrient | StemDiameterWeek4 | 0.019 | 0.181 |
| Prairie | High Nutrient | ChlorophyllWeek2 | -0.102 | -0.046 |
| Prairie | High Nutrient | ChlorophyllWeek4 | -0.334 | -0.027 |
| Prairie | High Nutrient | Emergence | -0.071 | -0.071 |
| Prairie | High Nutrient | HeightWeek2 | 0.098 | 0.139 |
| Prairie | High Nutrient | HeightWeek4 | 0.048 | 0.070 |
| Prairie | High Nutrient | RootBiomass | 0.077 | 0.135 |
| Prairie | High Nutrient | ShootBiomass | 0.078 | 0.090 |
| Prairie | High Nutrient | StemDiameterWeek2 | -0.129 | -0.030 |
| Prairie | High Nutrient | StemDiameterWeek4 | -0.154 | -0.037 |
| Prairie | Low Nutrient | ChlorophyllWeek2 | -0.121 | 0.049 |
| Prairie | Low Nutrient | ChlorophyllWeek4 | -0.011 | -0.005 |
| Prairie | Low Nutrient | Emergence | 0.154 | 0.200 |
| Prairie | Low Nutrient | HeightWeek2 | 0.087 | 0.099 |
| Prairie | Low Nutrient | HeightWeek4 | 0.103 | 0.141 |
| Prairie | Low Nutrient | RootBiomass | 0.165 | 0.244 |
| Prairie | Low Nutrient | ShootBiomass | 0.170 | 0.250 |
| Prairie | Low Nutrient | StemDiameterWeek2 | -0.127 | -0.034 |
| Prairie | Low Nutrient | StemDiameterWeek4 | -0.069 | 0.087 |

